# Evolutionary stasis of an RNA virus indicates arbovirus re-emergence triggered by accidental release

**DOI:** 10.1101/2019.12.11.872705

**Authors:** David J Pascall, Kyriaki Nomikou, Emmanuel Bréard, Stephan Zientara, Ana da Silva Filipe, Bernd Hoffmann, Maude Jacquot, Joshua B. Singer, Kris De Clercq, Anette Bøtner, Corinne Sailleau, Cyrille Viarouge, Carrie Batten, Giantonella Puggioni, Ciriaco Ligios, Giovanni Savini, Piet A. van Rijn, Peter PC Mertens, Roman Biek, Massimo Palmarini

**Affiliations:** Institute of Biodiversity, Animal Health and Comparative Medicine, Boyd Orr Centre for Population and Ecosystem Health, University of Glasgow, Glasgow, United Kingdom; MRC-University of Glasgow Centre for Virus Research, Glasgow, United Kingdom; The School of Veterinary Medicine and Science, University of Nottingham, Sutton Bonington, Leicestershire, United Kingdom; UMR Virologie, INRA, Ecole Nationale Vétérinaire d’Alfort, Laboratoire de Santé Animale d’Alfort, ANSES, Université Paris-Est, 94700 Maisons-Alfort, France; Institute of Diagnostic Virology, Friedrich-Loeffler-Institut, Greifswald-Insel Riems, Germany; Spatial Epidemiology Lab (SpELL), University of Brussels, Brussels, Belgium; INRAE-VetAgro Sup, UMR Epidemiology of Animal and Zoonotic Diseases, F-63122, Saint Genès-Champanelle, France; Infectious Diseases in Animals, Exotic and Particular Diseases, Sciensano, 1180 Brussels, Belgium; Section for Veterinary Clinical Microbiology, Department of Veterinary and Animal Sciences, University of Copenhagen, Denmark; Department of Virus & Microbiological Special Diagnostics, Statens Serum Institut, Copenhagen, Denmark; The Pirbright Institute, Ash Road, Pirbright, Woking, Surrey, GU24 0NF, United Kingdom; Istituto Zooprofilattico Sperimentale della Sardegna, Via Duca degli Abruzzi, 07100, Sassari, Italy; Istituto Zooprofilattico Sperimentale dell’Abruzzo e del Molise (IZSAM), Teramo, Italy; Department of Virology, Wageningen Bioveterinary Research (WBVR), Lelystad, The Netherlands; Department of Biochemistry, Centre for Human Metabolomics, North-West University, Potchefstroom, South Africa

## Abstract

The mechanisms underlying virus emergence are rarely well understood, making the appearance of outbreaks largely unpredictable. Bluetongue virus serotype 8 (BTV-8), an insect-borne virus of ruminants, emerged in livestock in Northern Europe in 2006, spreading to most European countries by 2009 and causing losses of billions of Euros. Though the outbreak was successfully controlled through vaccination by early 2010, puzzlingly a closely-related BTV-8 strain re-emerged in France in 2015, triggering a second outbreak that is still ongoing. The origin of this virus and the mechanisms underlying its re-emergence are unknown. Here, we performed phylogenetic analyses of 164 whole BTV-8 genomes sampled throughout the two outbreaks. We demonstrate consistent clock-like virus evolution during both epizootics but found negligible evolutionary change between them. We estimate that the ancestor of the second outbreak dates from the height of the first outbreak in 2008. This implies that the virus had not been replicating for multiple years prior to its re-emergence in 2015. Given the absence of any known natural mechanism that could explain BTV-8 persistence over this period without replication, we conclude that the second outbreak was most likely initiated by accidental exposure of livestock to frozen material contaminated with virus from approximately 2008. Our work highlights new targets for pathogen surveillance programmes in livestock and illustrates the power of genomic epidemiology to identify pathways of infectious disease emergence.

## Introduction

Infectious disease outbreaks are a major burden on human and animal health. They can dramatically reduce the productivity of entire countries, due to direct losses, control measures, trade bans or public fear [1]. Diseases caused by insect-borne viruses (arboviruses) in particular, have increased substantially in recent decades [2-4] and there is an urgent need to better understand the causes of their emergence in order to devise better control and prevention strategies. The factors leading to disease emergence are often unclear and case studies of intensely studied outbreaks can therefore provide important wider lessons.

Bluetongue is a major disease of domestic ruminants caused by the bluetongue virus (BTV), an arbovirus transmitted by *Culicoides* midges. BTV is the type species of the genus Orbivirus, within the family *Reoviridae* and possesses 10 double-stranded RNA genome segments encoding for 7 structural and 4 or 5 non-structural proteins [5-7]. BTV infection in sheep can induce a variety of clinical outcomes, which in the most extreme cases include a lethal hemorrhagic fever [8-10]. Infection in cows and goats results instead in milder and often sub-clinical clinical signs [8, 9]. BTV can also infect wild ruminants and, more rarely, other mammal species [11-15].

Like many other arboviruses, the geographical spread of BTV has increased significantly in the last 20 years [16-18]. In August 2006, BTV serotype 8 (BTV-8) emerged for the first time in the Netherlands [19-24], leading to dramatic losses of sheep and causing extensive economic damage to farming communities, costing on the order of billions of euros [25-28]. The virus quickly spread across the continent, with confirmed infections in 16 countries by 2008 (Fig 1). The outbreak was ultimately controlled through a pan-European vaccination campaign, using inactivated vaccines, with a few last cases detected in Europe in 2010 [29]. However, after a five-year period with no BTV-8 cases recorded throughout Europe, the virus re-emerged in France [30] and has since continued to spread. France was declared enzootic in 2018 and recent cases reported in adjacent countries, including Germany, Switzerland and Belgium [31].

**Figure 1.**
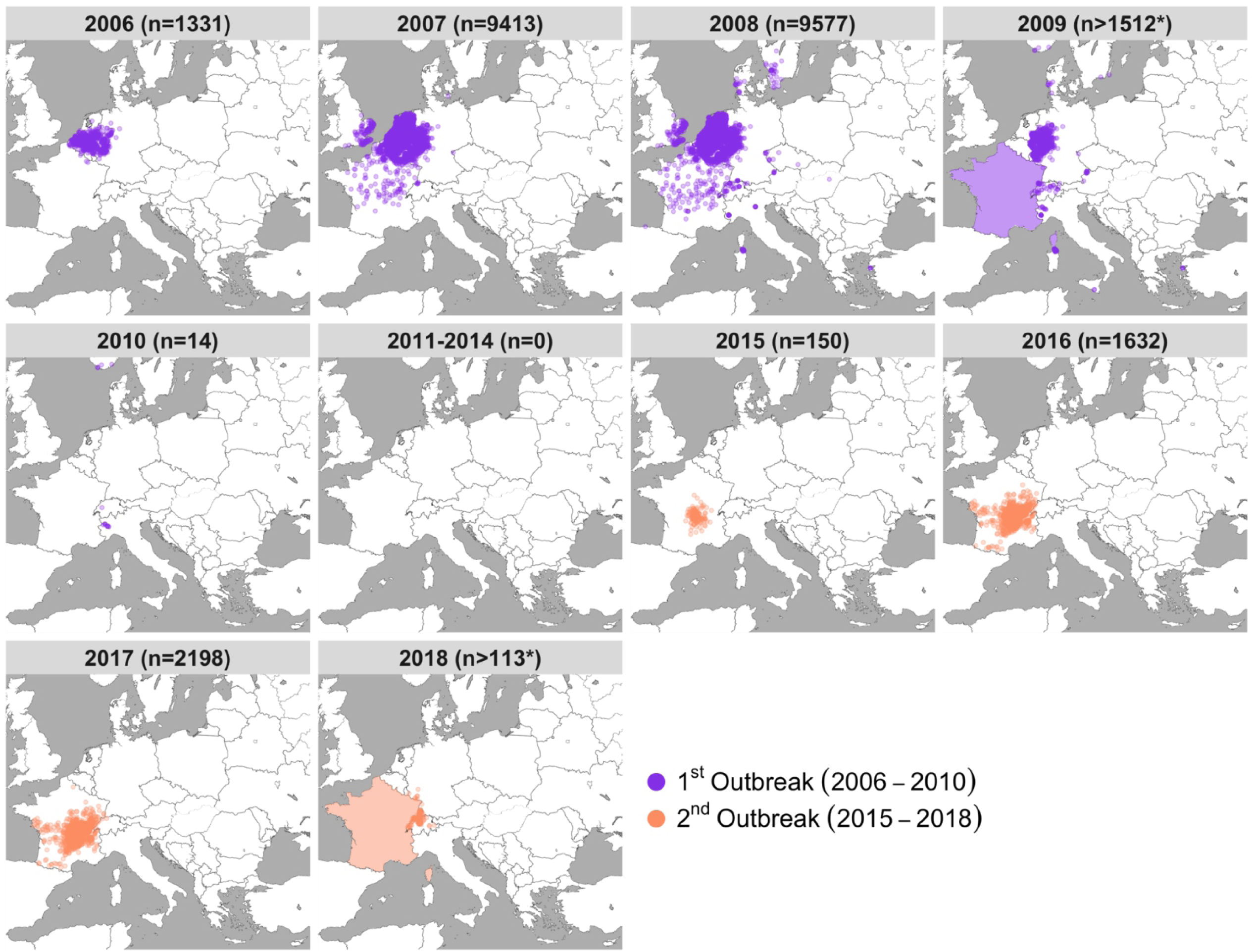
Emergence and re-emergence of BTV-8 in Europe. Location and number of premises housing livestock infected with Bluetongue virus serotype 8 collected from “immediate reports” to the OIE (World Organisation for Animal Health). Data was accessed from the WAHIS database on the 12^th^ of July 2019. Immediate report data was provided by the OIE. Each point corresponds to one infected premise, with total counts for each year (n), with the first outbreak shown in purple and the second outbreak shown in orange. Please note that maps includes only immediate notification to the OIE and this is therefore a smaller number than the totality of affected premises during the outbreaks. In addition, France is shaded in 2009 and 2018 as France stopped providing immediate reporting, requiring location data, to the OIE. As such, the counts of infected premises from 2009 and 2018 are not comparable to previous years as they exclude France, where the virus was widespread. Map adapted from tiles by Stamen Design, under Creative Commons (CC BY 3.0) using data by OpenStreetMap, under the Open Database Licence.

The source and mechanism of BTV-8 re-emergence in France remains obscure. Initial genetic data from one isolate, suggested the re-emerging virus in France to be a close relative of the lineage causing the 2006-2010 outbreak [30, 32, 33]. The prevailing theory was that the virus had continued to be transmitted sub-clinically but remained unrecorded in livestock or wild ruminants after it had been declared absent from Europe in 2011 [30]. However, there is currently little evidence to support this hypothesis. Based on serological evidence, wild ungulates do not appear to have sustained transmission [34, 35]. Similarly, serological testing of cattle sampled in 2014, indicated a rapid decline of seropositivity after vaccination ceased in France in 2010, consistent with a new (re-) introduction of the virus in, or just before, 2015 [34, 36, 37].

We describe the use of phylogenetic and evolutionary analyses of BTV-8 virus samples, collected during the first and second European outbreaks, to gain insights into the mechanisms that allowed BTV-8 to re-emerge in France in 2015. For this, we generated a novel data set of full genome sequences for more than 150 viruses sampled throughout both outbreaks. We show that the evolutionary signatures contained in these data are inconsistent with continuous circulation of BTV-8 between the outbreaks and instead point to re-emergence being caused by an anthropogenic factor, such as accidental release of contaminated material.

## Results

### Viruses from the first and second European BTV-8 outbreaks form a single monophyletic clade

We analysed newly sequenced full genomes of 153 BTV-8 samples collected from infected sheep and cattle throughout the BTV-8 outbreaks in Europe along with 11 BTV-8 isolates previously published. Samples from the first outbreak were collected from infected animals in 10 different countries between 2006 and 2009, while samples from the second outbreak were collected from France between 2015 and 2018 (Table S1). To minimise or exclude the possibility of including genome mutations acquired during extensive passage of the virus in culture, we sequenced the great majority of samples directly from clinical samples (blood) of infected animals or from isolates kept in culture for a minimum number of passages (Table S1).

A maximum likelihood (ML) tree revealed considerable genetic diversity both within the first (2006-2010) and the second outbreak (2015-2018) (Fig 2). The tree showed that all sequences from the second European outbreak form a well-supported monophyletic clade that is nested within the virus lineages circulating during the first outbreak in 2006-2010 (Fig 2, Fig S1). Specifically, the clade of the second outbreak derives from a clade from the first outbreak, including predominantly viruses from France and Germany collected in 2007 and 2008. The viruses from the second French outbreak can be distinguished into two further clades, one including viruses from 2015 and 2016, the other including samples spanning the entire outbreak (2015-2018). Interestingly, both clades were already present among the eight samples from the farm in France (in Auvergne-Allier) from which the first diagnosis of re-emerged BTV-8 was made in August/September of 2015. Surprisingly, the branch leading to the re-emerging virus appeared short, given the 5-year period between the outbreaks, implying a slow rate of evolution during this period (Fig 2).

**Figure 2.**
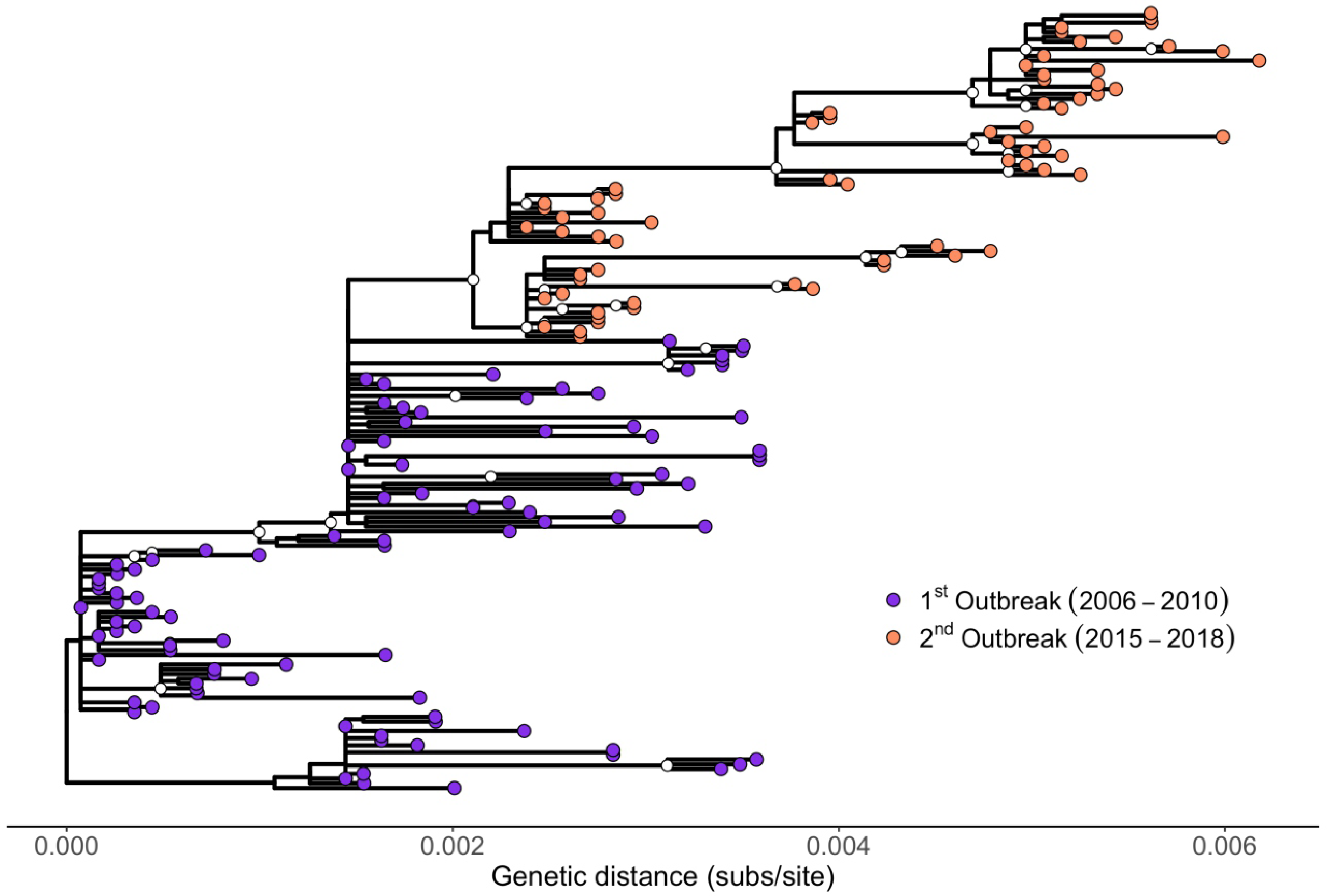
Phylogenetic tree of 164 BTV-8 samples collected during the European outbreak between 2006 and 2018. Maximum likelihood tree estimated in PhyML. The scale shows substitutions per site. Clades represented 700 or more times within 1000 bootstraps are indicated by a white circle. Samples from the first outbreak are shown with purple circles while samples form the second outbreak are shown with an orange circle. Note that an identical tree with labels corresponding to the individual samples is shown as Fig S1.

### BTV-8 re-emergence associated with exceptionally slow evolution

To test if the lower amount of divergence along the branch separating the two outbreaks was unusual, we estimated the evolutionary rate of BTV-8 from the set of 164 genomes. For this, we applied a lognormal relaxed clock model that allows for branch-specific heterogeneity in clock rates, implemented in the Bayesian phylogenetic software BEAST (Fig 3 and Fig S2). The mean evolutionary rate estimate was 4.04×10^−4^ substitutions per site per year (95% HPD: 3.37×10^−4^, 4.72×10^−4^), corresponding to an expected 7.76 substitutions per year (95% HPD: 6.47, 9.06) across the entire BTV-8 genome. In contrast, the emerging branch had an estimated mean evolutionary rate that was nearly an order of magnitude slower at 8.24×10^−5^ substitutions per site per year (95% HPD: 3.93×10^−5^, 1.32×10^−4^). Indeed, within the presented maximum clade credibility tree, this branch had the lowest median rate across the posterior distribution of trees.

**Figure 3.**
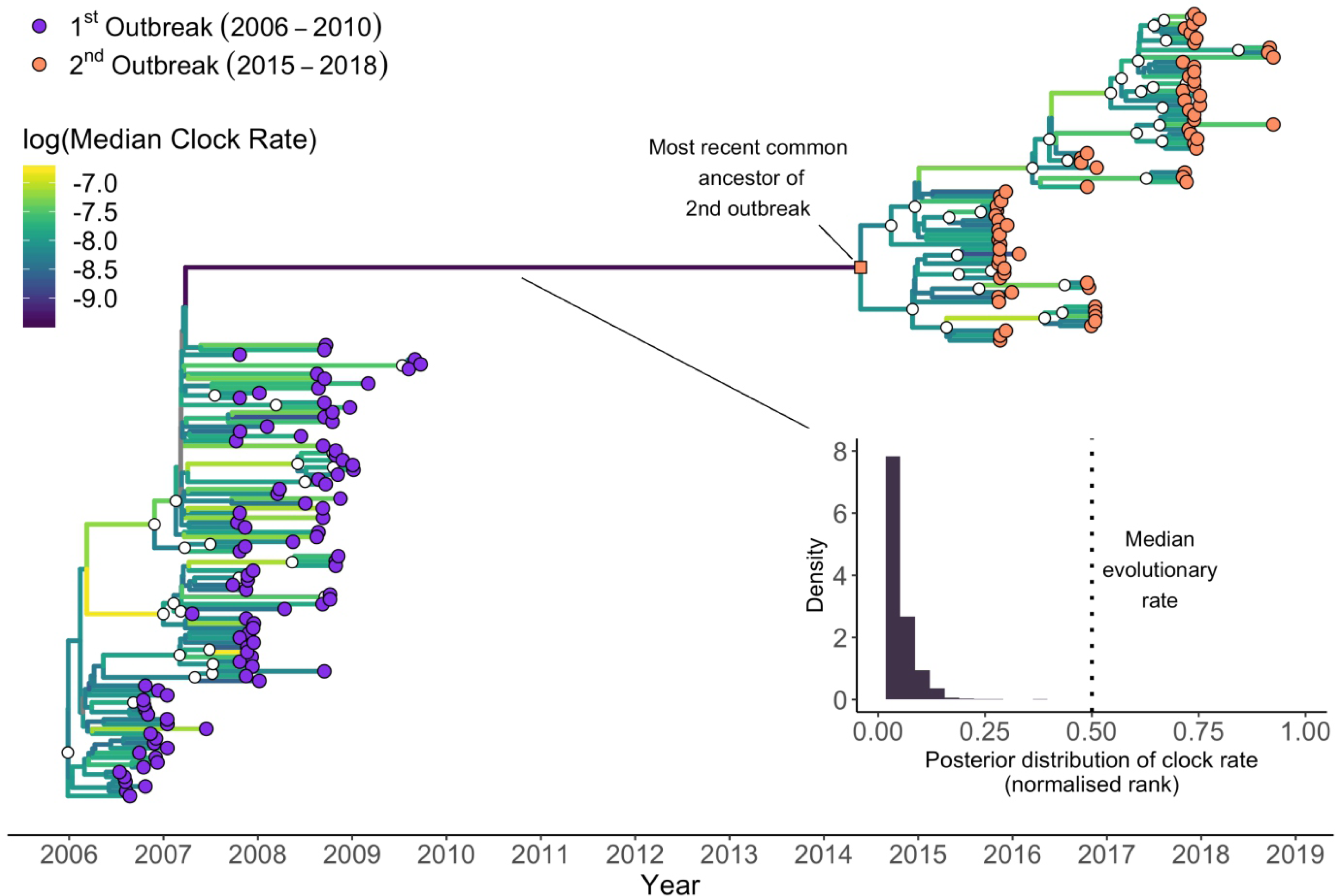
Time-scaled phylogenetic tree of BTV-8 samples collected during the European outbreaks between 2006 and 2018. Maximum clade credibility time-calibrated phylogenetic tree generated in BEAST. The tree is scaled in years, with the final sampling date being October 2018. Clades with posterior support of 0.9 or higher are indicated by a white circle. Samples from the first outbreak are shown with a purple circle while samples from the second are shown with orange circles. The branches are coloured accordingly to their median evolutionary rate across the posterior (see heatmap within the figure). The long branch leading from the first outbreak to the second (in dark purple) shows the slowest evolutionary rate on the maximum clade credibility tree. The inset shows the posterior distribution of the normalised rank of the evolutionary rate of the long branch relative the other branches in the tree, as estimated from the lognormal relaxed clock. Values close to 0 represent slow evolution relative to the rest of the tree, values close to 1 represent fast evolution relative to the rest of the tree, and 0.5 represents the median branch evolutionary rate on the tree. This analysis show an unusually slow evolution of BTV-8 between outbreaks. See the methods section for the specifics of this calculation. Note that an identical tree with labels corresponding to the individual samples is shown as Fig S2.

Using BEAST, we reconstructed the sequence of the most recent common ancestor of the viruses sequenced from the second outbreak. This ancestral sequence displayed only 7 nucleotide substitutions (of which 6 were synonymous or in the untranslated regions of the viral genome) compared to BTV-8_FRA2007-3673_, the genetically closest virus within the data set, which was collected from France in August 2007 (Fig 4A). In comparison, BTV-8_FRA2007-3673_ displayed 23 nucleotide substitutions (of which 16 were synonymous or in the UTR) compared to the first sequence available from the first outbreak and collected in August 2006 in the Netherlands (BTV-8_NET2006-04_) (Fig 4A). The number of mutations of BTV-8_FRA2007-3673_ compared to the BTV-8 sequences in the dataset in 2006 (n=23) ranges between 15 and 23 while those compared to the virus sequences collected in 2008 (n= 37) varies between 2 and 56. Hence, sequence variation between BTV-8 samples collected only a year apart during the first outbreak is in general far higher than that between the ancestor of the re-emerged BTV-8 strain and its closest relative in the first outbreak.

**Figure 4.**
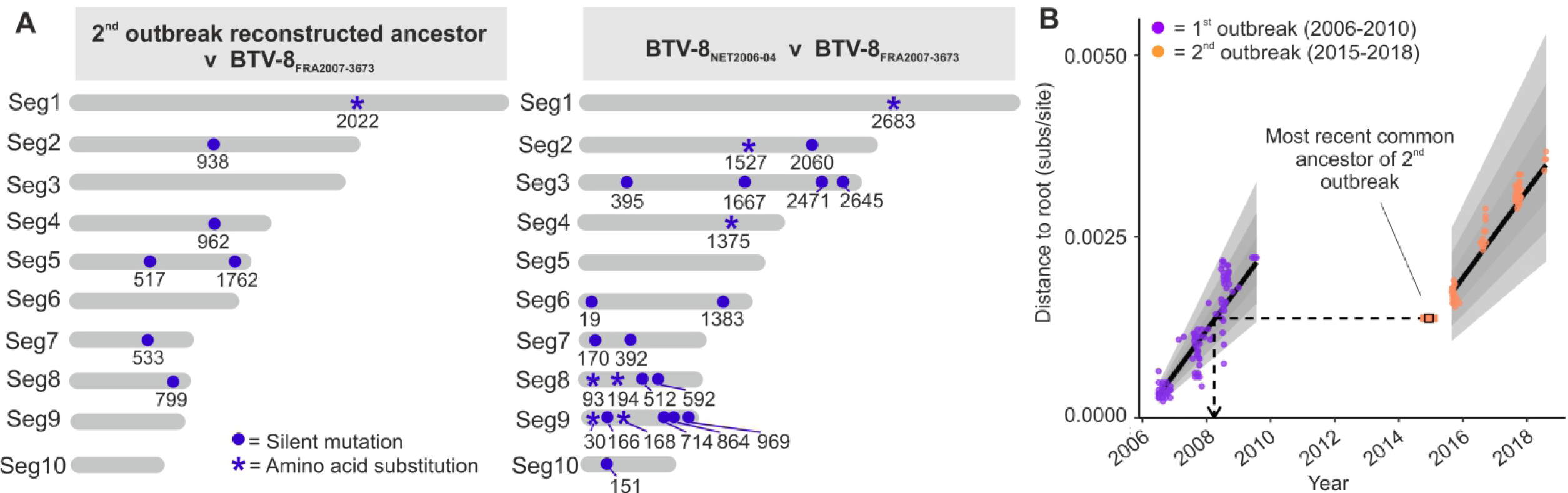
Lack of evolution of BTV-8 between the two European outbreaks. A. Graphic representation of nucleotide substitutions between the genomes of the earliest BTV-8 collected from the 1^st^ European outbreak (BTV-8_NET2006-04_), the reconstructed ancestor of the second BTV-8 outbreak (BTV-8_FRA2015_) and the most similar virus to the latter sequence present in our dataset (BTV-8_NET2007-3673_). Substitutions are shown as a blue circle with numbers indicating the genomic position for each of the 10 genomic segments. Asterisks indicate those mutations inducing also an amino acid substitution. B. Genetic divergence of 164 BTV-8 samples collected from the two European outbreaks against their sampling date (circles). The regression lines from the best fitting linear model for each outbreak are shown in black. Error regions around the lines correspond to 50%, 80% and 95% posterior predictive intervals. When the day was unknown, the date was fixed to the 16th of the month, and when the month was unknown, the date was fixed to the midpoint of the year. The inferred age of the ancestor of the second outbreak is shown in blue, with a 95% highest posterior density error. The dashed line indicates that a virus of the degree of divergence inferred for the ancestor of the second outbreak is consistent with virus from the first outbreak circulating in 2008.

### “Frozen evolution”: clock-like evolution of BTV-8 during, but not in-between, outbreaks

Next, we included the reconstructed ancestral sequence in our dataset and re-estimated the ML tree to get measures of genetic distance from the root of the tree. Consistent with clock-like evolution, the genetic distance between virus sample and the root increased linearly with time during both the first (TempEst: slope=7.2708×10^−4^ subs/site/year, r^2^=0.8113) and second outbreaks (TempEst: slope = 6.9291×10^−4^, r^2^= 0.9605). There was no evidence that the evolutionary rate of the virus differed between the two outbreaks (p-value for Date:Outbreak interaction = 0.194). However, there is a clear discontinuity in the accumulation of mutations between the two outbreaks, consistent with a period where clock-like evolution had essentially ceased. Consequently, the reconstructed sequence of the ancestor of the second outbreak, when included in the ML tree, has an inferred distance from the tree root that is consistent with a virus from late March 2008 according to the root to tip regression (Fig 4B).

## Discussion

Diseases of livestock can be exceedingly interesting models to study virus emergence, given that harmonised international surveillance systems and regulatory frameworks provide opportunities to access field samples with associated metadata across national borders. Here, the BTV-8 European outbreaks provided us with the opportunity to investigate the mechanisms surrounding arbovirus emergence based on a uniquely rich dataset. Our results indicate that the re-emergence of BTV-8 in France in 2015 was caused by a virus that exhibits a lack of evolutionary changes since the first outbreak. This is inconsistent with the prevalent view of undetected low-level circulation of the virus in wild or domestic ruminants, between 2010 and 2015, and instead points to another mechanism of emergence.

We showed a large discontinuity in the number of mutations accumulated by BTV-8 between 2010 and 2015, even though the evolutionary rates of the virus during the first and second outbreak were indistinguishable and of the same order as rates reported in previous BTV studies [38, 39]. If the virus had been replicating consistently in an undetected population from 2010 to 2015, we would expect the genetic distance of the isolates from the second European outbreak to continue the trend of increased divergence after the first outbreak. However, the sequences from the second outbreak exhibit genetic divergences that fall considerably below what would be expected if the trend-line from the first outbreak was extended, illustrating a paucity of mutations relative to expectations (Fig 4B). Indeed, the divergence of the reconstructed ancestor of the second bluetongue outbreak is consistent with the virus stopping replication in March 2008. The lack of divergence is also illustrated by the fact that the reconstructed ancestor of the BTV-8 outbreak has only 7 mutations separating it from its closest relative in the analysed dataset, a French sample collected in August 2007 (BTV-8_FRA2007-3673_), despite putatively having been replicating for at least half a decade after that samples’ collection. In comparison, BTV-8_FRA2007-3673_ showed 23 mutations compared to the genome of the earliest BTV-8 sample obtained from the Netherlands in August 2006, only a year earlier. The corresponding rate of evolution estimated for the emerging branch was almost an order of magnitude slower than the mean clock rate, highlighting it as exceptionally slow (Fig 3). Moreover, we hypothesise that some or all of the estimated seven mutations on this branch might have been accumulated during the first outbreak, given that the emerging branch connects to an internal node in the time-scaled phylogeny with a date of early 2007, at the height of the first outbreak. The subsequent accumulation of seven mutations is consistent with the idea that this virus continued to circulate until early 2008 (the inferred date from the root-to-tip regression) and then ceased to change all together until its re-emergence in 2015.

Given the unexpectedly low number of mutations observed between the two outbreaks, our data indicate that the common ancestor of the second European outbreak either ceased, or dramatically slowed its replication, in early 2008. This is inconsistent with current knowledge of the biology of BTV and RNA viruses in general. For example, a potential explanation could be that BTV persistently infected a host for several (5 to 8) years, with little or no replication, before being re-activated and starting the second outbreak. While this may be possible with DNA viruses, or RNA viruses with a DNA intermediate [40-45], it has never been described for Reoviruses such as BTV or other RNA viruses. Rabies virus may provide an exception based on a handful of case reports of virus reactivation after latency of several years [46] but it has not been documented whether these involved a lack of evolutionary changes. In other cases, as in foot-and-mouth disease, viral RNA and infectious virus have been shown to persist in reservoir hosts for multiple years. However, re-isolation of virus (as opposed to detection of viral RNA) indicates that the virus replicates during persistent infection and accumulates nucleotide substitutions at a rate comparable to actively replicating viruses [47, 48]. Hepatitis C virus for example is also known to persist in a number of patients for a number of years but, again, with continuing viremia, and thus virus replication [49].

Another hypothetical scenario could be envisaged if BTV-8 was to remain “latent” in midges’ eggs for a number of years. However, there is no evidence of vertical transmission in BTV-infected Culicoides [50-53]. This, in conjunction with the need for infected midge eggs to survive for years, rather than a single overwintering season, makes this scenario highly unlikely.

Overall, we judge the possibility of persistence of BTV-8 in a mammalian or invertebrate host for longer than 5 years, in the absence of viral replication, followed by viral reactivation and subsequent onwards spread, to be implausible. Our data instead point to the release of a BTV-8 virus retained from the initial outbreak by other means as the cause of the second outbreak in France in 2015. Anthropogenic causes of virus outbreaks have been described before. Accidental virus release is thought to have been responsible for the 1977 influenza A H1N1 outbreak, caused by a virus which closely matched a variant circulating in the 1950s [54]. Likewise, the 1995 Venezuelan equine encephalitis subtype IC epidemic which was caused by a virus closely related to a strain circulating in 1962-64 [55]. For livestock pathogens, a localised outbreak of FMDV in the UK in 2007 was linked to virus escaped from research facilities [56].

Our data cannot reveal the anthropogenic source from which BTV-8 was re-introduced in France in 2015. We speculate that laboratory escape of virus preparations, such as the case of FMDV in the UK in 2007, is unlikely as BTV needs an insect vector for efficient transmission and we are unaware of any *in-vivo* insect experiments in France with BTV during that period. However, due to specific animal husbandry procedures, there are important potential sources of frozen virus that apply to livestock viruses that are not present in viruses of most other animals, specifically the widespread use of bull semen for artificial insemination and embryo transfer in cows [57, 58]. BTV has been detected in the semen of viremic bulls and rams, can initiate infection in the mother and be transmitted vertically to the embryo [59, 60]. Additionally, contaminated embryos can cause transmission on implantation [61]. As such, both semen and embryos may represent potential sources of BTV infection. Contaminated frozen colostrum may also be a potential source, considering that oral transmission has been shown to be possible with BTV-8 [62]. However, it is not normal practice to keep colostrum frozen for a number of years. Interestingly, while international regulations specify that bull donors and semen that are exported internationally must be screened for various pathogens including BTV [63], this does not apply to premises trading only locally and carrying out private insemination procedures [64]. Thus, semen from a BTV-8 infected bull could have been collected or an embryo generated from an infected but asymptomatic animal and used years later without detection. The eventual use of this frozen animal product could have then led to the infection that caused the second Western European outbreak.

We stress that the link between bull semen trade and embryo implantation in France and the BTV-8 re-emergence in 2015 is only speculative. However, we have shown that the re-emergence of BTV-8 in France in 2015 is unlikely to be due to cryptic continuing transmission and we can exclude a reintroduction from another endemic country. Thus, our data are incompatible with the two current dominant theories for explaining the 2015 outbreak [30]. The lack of accumulated mutations in the virus implies that there was either an ongoing persistent infection in the absence of viral replication for several years, or the virus was reintroduced by humans from material that had been frozen during the first outbreak. We argued the second of these explanations to be more likely. Our findings highlight new areas requiring thorough surveillance programmes for the control of infectious disease of livestock. In addition, our approach illustrates how unrecognised pathways of disease emergence can be revealed using pathogen genomic epidemiology.

## Methods

### Samples

Blood samples from animals infected with BTV-8 were received from 10 European countries during the bluetongue outbreaks from 2006 to 2018. In some instances, samples analysed were viruses isolated in tissue culture from blood of infected animals. Table S1 provides the metadata related to the dataset used in this study. These include virus strain names, animal species of origin, geographical location and date of sampling. In addition, metadata include whether the viral genome sequence was obtained directly from clinical material (blood) or from an isolate in tissue culture, sequencing methods and GenBank accession number.

### RNA extraction and Illumina library preparation

Total RNA was extracted from infected blood samples, and virus isolates using Trizol LS^®^ (Invitrogen, USA) and purified using Direct-zol RNA MiniPrep (Zymo Research, USA) as per manufacturer’s protocol. RNA samples were treated with DNAse I (Ambion) and purified with 3X Agencourt RNAClean XP beads (Beckman Coulter, USA). Total RNA concentration was quantified using the Qubit Fluorimeter (Life Technologies, USA) and Qubit RNA HS Assay (Life Technologies, USA), while RNA integrity was assessed using Agilent 4200 TapeStation (Agilent, USA). In order to avoid cross-contaminations, RNA extractions, from virus isolates was performed separately from those of infected blood samples. Similarly, RNA extractions were carried out separately on the bases of geographical origin and year of collection of the samples. In addition, library preparations, target enrichment and sequencing runs (see below) were carried out also on separate days following the same criteria than above. Libraries from low and without measurable RNA (low input) were also prepared separately from those with measurable RNA (high input). Libraries were prepared for Illumina sequencing using the Illumina TruSeq Stranded mRNA HT kit (Illumina) using 5 μl of sample RNA (up to 250 ng of total RNA) according to the manufacturer’s instructions. Briefly, after RNA was fragmented, it was reverse transcribed using SuperScript II Reverse Transcriptase (Invitrogen, USA) and random hexamers. Single strand cDNA was immediately converted to double stranded cDNA, cleaned-up with Agencourt^®^ AMPure^®^ XP magnetic beads (Beckman Coulter, USA), quantified using Qubit Fluorimeter and Qubit dsDNA HS Assay Kit (Life Technologies, USA) and size distribution was assessed using a 4200 TapeStation System with High Sensitivity D1000 Screen Tape assay (Agilent, USA). A-tailing was performed followed by adapters ligation. After a purification step, dual indexed libraries were PCR amplified and the purified PCR products were pooled in equimolar concentrations and sequenced using 150 paired end sequencing on MiSeq or NextSeq500 sequencers (Illumina USA).

### Targeted enrichment sequencing

We carried out multiplexed viral targeted enrichment followed by Illumina sequencing using the NimbleGen SeqCap EZ system (Roche, USA), for improved viral detection from clinical material. This approach was followed in order to increase the number of BTV-8 samples from which we could obtain a complete viral genome sequence directly from clinical material, including those with very low amount of viral RNA. Libraries were prepared following the above described standard Illumina TruSeq Stranded mRNA protocol. They were quantified using Qubit Fluorometer and Qubit dsDNA high sensitivity (HS) Assay Kit (Life Technologies, USA). Quality and size distribution was validated using the High Sensitivity D1000 Screen Tape assay (Agilent) in a 4200 TapeStation System (Agilent, USA) and were normalized according to BTV viral load and mass. A 1000 ng aliquot of the pooled library was enriched using SeqCap EZ Developer Probes (Roche/NimbleGen) (see below), according to the manufacturer’s protocol. After a 14 cycle post enrichment PCR amplification, the cleaned PCR products were pooled and were sequenced with a 151-base paired-end reads on a NexSeq500, 550 cartridge (Illumina, USA). Probes were designed using all BTV sequences available on NCBI Genbank, RefSeq, DNA Data Bank of Japan (DDBJ) and EMBL EBI databases (as accessed by October 2016). The resulting NimbleGen biotinylated soluble capture probe library (“BTV-Cap”), contains a probe set of more than 500,000 probes, designed to minimise capture of *Culicoides sonorensis, Bos taurus, Ovis aries, Capra hircus* and *Mesocricetus auratus* genomes.

### Consensus calling

All consensus calling was performed on a cluster running Ubuntu v. 14.04.5 LTS. In all cases, BAM and SAM files were handled using samtools v. 1.3 [65]. The R packages ggplot2 [66], seqinr [67], stringr [68], and vcfR [69] were all used in scripts at various points in the following section. Paired end raw reads were trimmed with Trim Galore! v. 0.4.0 (http://www.bioinformatics.babraham.ac.uk/projects/trim_galore/) with a quality cut-off of 30. Any reads below 50 bp were discarded. Following this, any reads that were unpaired were discarded if they were under 100 bp. Overlapping reads were combined using FLASH v. 1.2.11 [70]. Reads were mapped using bowtie2 v. 2.3.4.2 [71] to a reference database containing all the segments of all the described strains of BTV in order to manually check for mixed infections. Read were allowed to have as many valid maps as could be found. Mapping statistics were generated with weeSAM v. 1.5 (https://github.com/centre-for-virus-research/weeSAM) and transcripts per million for each target were then generated using eXpress v. 1.5.1 [72]. Separately, for the consensus generation, reads were mapped using Tanoti v. 9th July 2018 (https://github.com/vbsreenu/Tanoti/tree/master/src) against a reference BTV-8 genome from the European BTV-8 outbreak (GenBank accession numbers: JX680447-JX680456). Different software was used for the quality control and consensus building steps as bowtie2 generates metadata required for the downstream quality control steps which Tanoti does not.

Variants from the BTV-8 reference were then called from the Tanoti alignment with lofreq* v. 2.1.2 [73] with the minimum coverage of the filtering step set to 5 and all other parameters at their default values. Any variants from the reference with an allele frequency of greater than 0.5 were replaced into their positions in the reference to build a new reference sequence. Reads were then remapped to this new reference. This process was then repeated either 5 times or until the reference generated after the process was identical to the reference at the start of the last round of mapping. Reads were then mapped again against this new reference and ambiguities were called. A base was called unambiguously if the allele frequency of the dominant allele was greater than 0.75, otherwise the base was called ambiguously over all alleles with a frequency of greater than 0.05 using a bespoke script in R (Data S1). For both of these consensus sequences, positions were masked with “N”s if the coverage at the site was less than 5 separate paired end reads.

Final sequences were processed and annotated for submission to GenBank using an extension to the BTV-GLUE resource (http://btv-glue.cvr.gla.ac.uk). GenBank accession numbers for each of the sequences in our dataset are indicated in Table S1.

### Quality control

A sequence showing evidence of mixed infection or contamination was discarded. Potential contamination and/or mixed infection were detected by finding sequences that met the following two criteria: i) visible numbers of reads mapping to serotypes other than BTV-8 or the closely related BTV-18 in segments 2 and 6; and ii) the presence of regions, which when aligned showed large numbers of unique SNPs and ambiguous nucleotides. In total, 8 samples were discarded due to mixed infection with different BTV strains and/or contamination. During quality control, segment 7 from the sample FRA2008-28 was also removed, as it represented an obvious reassortment from a distinct BTV strain, but the rest of the sample was preserved. We used GiRaF v 1.02 [74] and MrBayes v 3.2.7a [75] on all the samples to test for the presence of less obvious reassortments between serotypes. Within the GiRaF algorithm, per segment trees were run for 1000000 iterations, with 500000 iterations discarded as burn-in. All other parameters were left at their default values. No reassortment was detected, so we opted to use all segments in a single concatenated phylogenetic tree. However, it should be noted that, as there is little variation in many segments, our ability to detect reassortment between two distinct but phylogenetically related strains is correspondingly low.

### Phylogenetics

Two separate phylogenetic analyses were performed, a maximum likelihood analysis performed in PhyML v. 20120412 [76] and a Bayesian analysis performed in BEAST v. 1.10 [77].

### Maximum likelihood analysis

To explore the diversity of the outbreaks, we generated a maximum likelihood tree. All segments were concatenated into a single sequence and a phylogeny using the GTR+G+I nucleotide model was run in PhyML v. 20120412 [78]. All parameters were optimised by maximum likelihood. The algorithm was the best of NNI and SPR moves with 10 random starts with 1000 bootstraps being performed on the best tree found. A maximum likelihood tree containing all sequences was then run, using the same settings as the first, and 1000 bootstraps were performed on the best tree found in those 12 starts. Given the observed short branch between the first and second outbreaks, this tree was then rooted at the optimal root found from the tree containing only the sequences from the first outbreak generated under the same settings, as calculated by TempEst v. 1.5.1 [79]. The TempEst rooting procedure also confirmed clock-like evolution for this dataset.

### BEAST analysis

Using known break points, the sequence for each segment was split into the untranslated region, and the first, second and third codon positions of the coding sequence. In the 9th and 10th segments there are regions with overlapping open reading frames, these were also placed together in their own partition. Separate evolutionary models, linked across segments, were applied to each of these partitions. The segments shared a lognormal relaxed molecular clock [80]. Given the difficulty, caused by combinatorial explosion, of model selection when there are multiple partitions, we performed a pre-analysis model selection protocol. Each segment was concatenated and a model was chosen for each partition using jModelTest v 2.1.10 [81]. The best model by Akaike Infomration Criterion corrected for small sample size (AICc) that was implemented in BEAST 1.10 was used. This was a GTR model for the first codon position, a HKY for the second, GTR+G with 4 gamma categories for the third, K80 for the UTR and JC for the regions with overlapping ORFs. We used a GRMF skyride model [82] for the tree prior. When the sampling date was not exactly known, the age of the tip was estimated in the MCMC with a uniform prior over the period of uncertainty. All priors were left at their default values except for the mean of the lognormal distribution for the relaxed molecular clock which was given a lognormal(−7.6, 3) prior. In all cases ambiguous nucleotides were used in the tree likelihood. Two trees were run, one containing only the sequences from the 2015 outbreak, and one containing all sequences. The tree containing all sequences was used to reconstruct the sequence of the ancestor of all the viruses in the second outbreak. BEAST will reconstruct sequence even in locations where the majority of sequences show gaps in the alignment. As such, there were three nucleotides that we removed from the final reconstructed sequence corresponding to locations in the original multiple sequence alignment where all sequences but one had gaps. The BEAST XMLs for the two trees described above are available as Data S2.

### Downstream statistical analysis and figure generation

Observed genetic distance from the full ML tree and sampling date were combined in R. When the exact sampling date was unknown, if the day within the sampling month was unknown, the date was fixed to the 16th of the month, and when the month was unknown, the date was fixed to the midpoint of the year. General linear models were fitted to test if the evolutionary rate of the virus was the same between two outbreak in the glm function in base R. The models used a gamma distribution with an identity link. The regression equation for the first model was: Genetic distance from root ∼ Date + Outbreak. The regression equation for the second model was: Genetic distance from root ∼ Date + Outbreak + Date:Outbreak. After no evidence of was found of differential rates between the two outbreaks, the general linear model without the interaction was run in brms [83] to generate predictive intervals for Figure 4b. A normal(0, 10) prior was placed over the intercept, standard normal priors were placed over all regression coefficients and a gamma(0.01, 0.01) prior over the shape parameter. The normalised rank of the evolutionary rate of the long branch was calculated by; for each tree in the posterior, ranking the estimated evolutionary rate of each branch from slowest to fastest, extracting the rank for the long branch, subtracting 1 so that the minimum was 0, [83]then dividing by the number of branches minus 1, so that a number between 0 and 1 was generated. Figures used the following R packages: ggplot2 [66], ggtree [84], ggthemes [85], cowplot [86, 87], ggmap [87], viridis [88], tidybayes [89], lubridate [90], sp [91], raster [92], maptools [93], rgeos [94], rgdal [95], sf [96] and PBSmapping [97].

## Acknowledgement

This work was supported by the EU H2020 PALE-Blu grant (project No: 727393-2) and by the Wellcome Trust (092806/B/10/Z; 206369/Z/17/Z). The Quadro P6000 graphics card used in this research was donated by the NVIDIA Corporation. We are extremely grateful to the OIE for providing the data included in Fig 1. Results of the study are the sole responsibility of the authors and not the OIE. We are grateful to members of our laboratories for helpful comments to the manuscript.

---

**S1 Figure.**
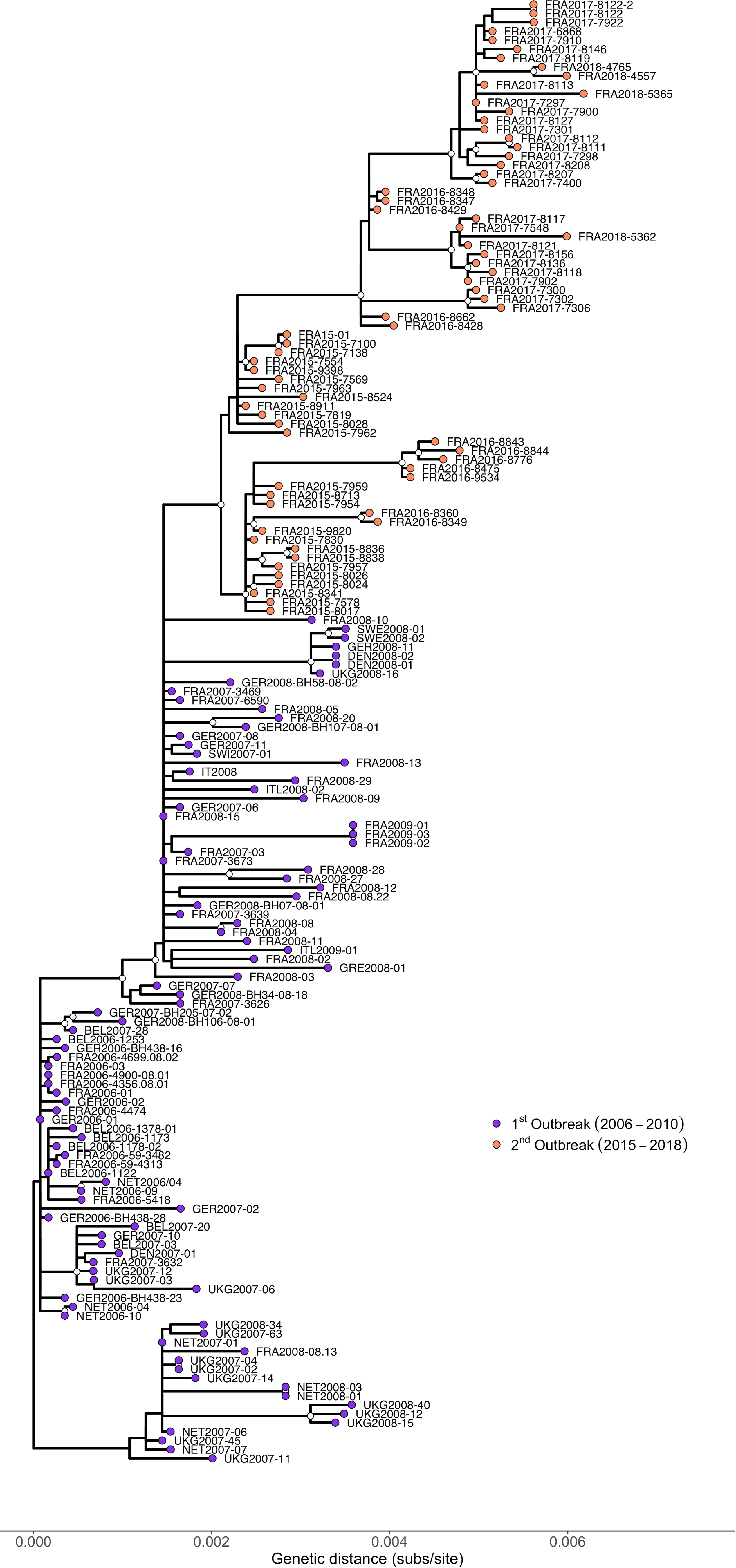

**Fig S2.**
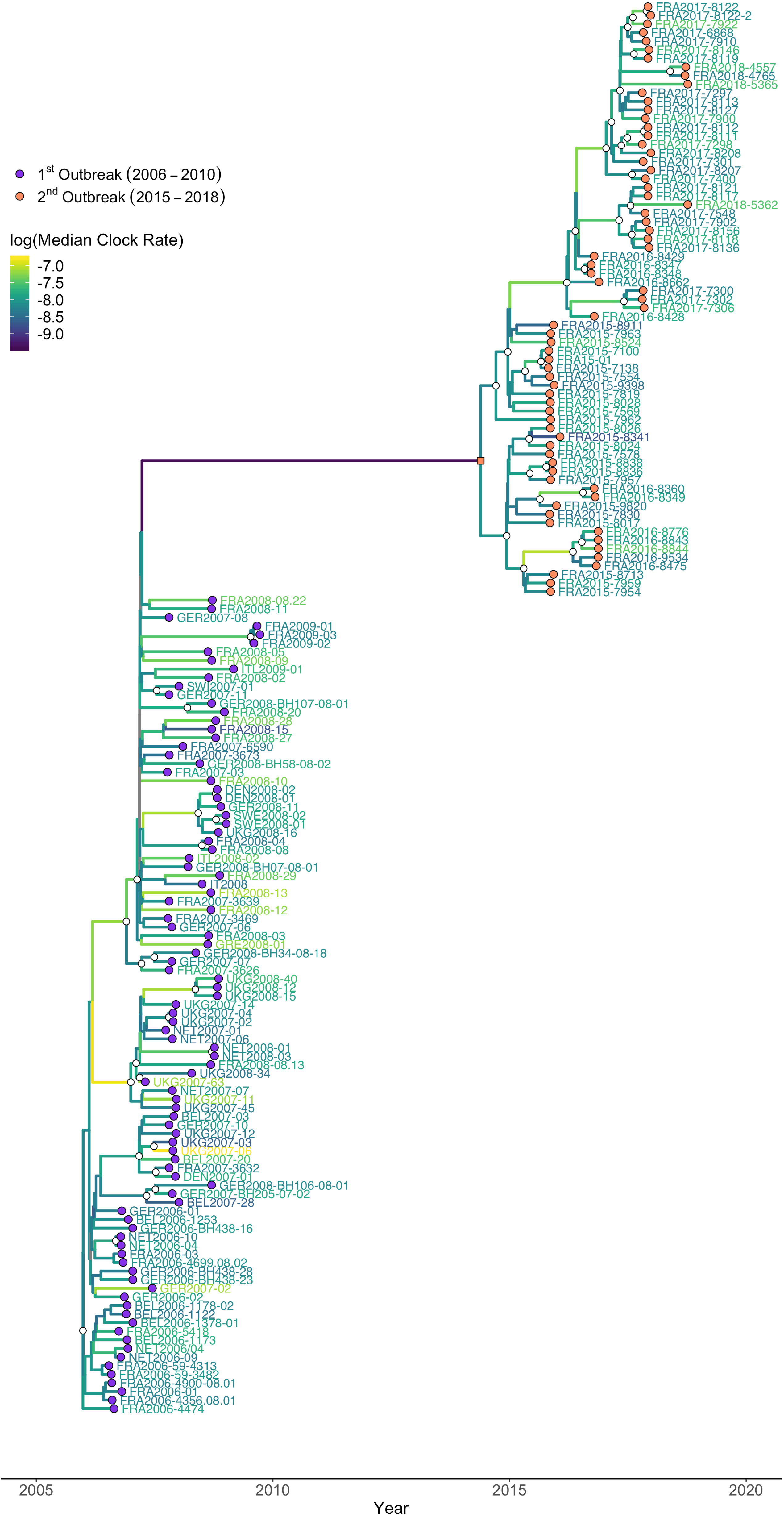

## References

1. Smith KM, Machalaba CC, Seifman R, Feferholtz Y, Karesh WB. Infectious disease and economics: The case for considering multi-sectoral impacts. One Health. 2019;7:100080.

2. Wilder-Smith A, Gubler DJ, Weaver SC, Monath TP, Heymann DL, Scott TW. Epidemic arboviral diseases: priorities for research and public health. The Lancet Infectious Diseases. 2017;17:e101–e6. doi: 10.1016/S1473-3099(16)30518-7.

3. Mayer SV, Tesh RB, Vasilakis N. The emergence of arthropod-borne viral diseases: A global prospective on dengue, chikungunya and zika fevers. Acta Trop. 2017;166:155–63.

4. Weaver SC, Reisen WK. Present and future arboviral threats. Antiviral Res. 2010;85(2):328–45.

5. Ratinier M, Caporale M, Golder M, Franzoni G, Allan K, Nunes SF, et al. Identification and characterization of a novel non-structural protein of bluetongue virus. PLoS Pathog. 2011;7(12):e1002477.

6. Mellor PS, Baylis M, Mertens PP. Bluetongue. Pastoret P-P, editor. London: Academic Press; 2009.

7. Roy P. Bluetongue virus: dissection of the polymerase complex. J Gen Virol. 2008;89(Pt 8):1789–804.

8. Caporale M, Di Gialleonorado L, Janowicz A, Wilkie G, Shaw A, Savini G, et al. Virus and host factors affecting the clinical outcome of bluetongue virus infection. J Virol. 2014;88(18):10399–411.

9. Maclachlan NJ, Drew CP, Darpel KE, Worwa G. The pathology and pathogenesis of bluetongue. J Comp Pathol. 2009;141(1):1–16.

10. Schwartz-Cornil I, Mertens PP, Contreras V, Hemati B, Pascale F, Breard E, et al. Bluetongue virus: virology, pathogenesis and immunity. Vet Res. 2008;39(5):46.

11. Clarke LL, Ruder MG, Kienzle-Dean C, Carter D, Stallknecht D, Howerth EW. Experimental Infection of White-Tailed Deer (Odocoileus Virginianus) with Bluetongue Virus Serotype 3. J Wildl Dis. 2019;55(3):627–36.

12. Linden A, Gregoire F, Nahayo A, Hanrez D, Mousset B, Massart AL, et al. Bluetongue virus in wild deer, Belgium, 2005-2008. Emerg Infect Dis. 2010;16(5):833–6.

13. Lopez-Olvera JR, Falconi C, Fernandez-Pacheco P, Fernandez-Pinero J, Sanchez MA, Palma A, et al. Experimental infection of European red deer (Cervus elaphus) with bluetongue virus serotypes 1 and 8. Vet Microbiol. 2010;145(1-2):148–52.

14. Mauroy A, Guyot H, De Clercq K, Cassart D, Thiry E, Saegerman C. Bluetongue in captive yaks. Emerg Infect Dis. 2008;14(4):675–6.

15. Jauniaux TP, De Clercq KE, Cassart DE, Kennedy S, Vandenbussche FE, Vandemeulebroucke EL, et al. Bluetongue in Eurasian lynx. Emerg Infect Dis. 2008;14(9):1496–8.

16. Wilson AJ, Mellor PS. Bluetongue in Europe: past, present and future. Philos Trans R Soc Lond B Biol Sci. 2009;364(1530):2669–81.

17. Baylis M, Caminade C, Turner J, Jones AE. The role of climate change in a developing threat: the case of bluetongue in Europe. Rev Sci Tech. 2017;36(2):467–78.

18. Maclachlan NJ, Zientara S, Wilson WC, Richt JA, Savini G. Bluetongue and epizootic hemorrhagic disease viruses: recent developments with these globally re-emerging arboviral infections of ruminants. Curr Opin Virol. 2019;34:56–62.

19. Elbers AR, Popma J, Oosterwolde S, van Rijn PA, Vellema P, van Rooij EM. A cross-sectional study to determine the seroprevalence of bluetongue virus serotype 8 in sheep and goats in 2006 and 2007 in the Netherlands. BMC Vet Res. 2008;4:33.

20. Mehlhorn H, Walldorf V, Klimpel S, Jahn B, Jaeger F, Eschweiler J, et al. First occurrence of Culicoides obsoletus-transmitted Bluetongue virus epidemic in Central Europe. Parasitol Res. 2007;101(1):219–28.

21. Szmaragd C, Wilson A, Carpenter S, Mertens PP, Mellor PS, Gubbins S. Mortality and case fatality during the recurrence of BTV-8 in northern Europe in 2007. Vet Rec. 2007;161(16):571–2.

22. Elbers AR, Backx A, Ekker HM, van der Spek AN, van Rijn PA. Performance of clinical signs to detect bluetongue virus serotype 8 outbreaks in cattle and sheep during the 2006-epidemic in The Netherlands. Vet Microbiol. 2008;129(1-2):156–62.

23. Elbers AR, Backx A, Meroc E, Gerbier G, Staubach C, Hendrickx G, et al. Field observations during the bluetongue serotype 8 epidemic in 2006. I. Detection of first outbreaks and clinical signs in sheep and cattle in Belgium, France and the Netherlands. Prev Vet Med. 2008;87(1-2):21–30.

24. Elbers AR, Backx A, Mintiens K, Gerbier G, Staubach C, Hendrickx G, et al. Field observations during the Bluetongue serotype 8 epidemic in 2006. II. Morbidity and mortality rate, case fatality and clinical recovery in sheep and cattle in the Netherlands. Prev Vet Med. 2008;87(1-2):31–40.

25. Velthuis AG, Saatkamp HW, Mourits MC, de Koeijer AA, Elbers AR. Financial consequences of the Dutch bluetongue serotype 8 epidemics of 2006 and 2007. Prev Vet Med. 2010;93(4):294–304.

26. Rushton J, Lyons N. Economic impact of Bluetongue: a review of the effects on production. Vet Ital. 2015;51(4):401–6.

27. Hasler B, Howe KS, Di Labio E, Schwermer H, Stark KD. Economic evaluation of the surveillance and intervention programme for bluetongue virus serotype 8 in Switzerland. Prev Vet Med. 2012;103(2-3):93–111.

28. Pinior B, Lebl K, Firth C, Rubel F, Fuchs R, Stockreiter S, et al. Cost analysis of bluetongue virus serotype 8 surveillance and vaccination programmes in Austria from 2005 to 2013. Vet J. 2015;206(2):154–60.

29. Zientara S, Sanchez-Vizcaino JM. Control of bluetongue in Europe. Vet Microbiol. 2013;165(1-2):33–7.

30. Sailleau C, Bréard E, Viarouge C, Vitour D, Romey A, Garnier A, et al. Re-Emergence of Bluetongue Virus Serotype 8 in France, 2015. Transboundary and emerging diseases. 2017;64:998–1000.

31. Gale P, Gauntlett F, Bowen J. Bluetongue virus (BTV-8) in Germany and Belgium. Department for Environment, Food and Rural Affairs, 2019.

32. Courtejoie N, Durand B, Bournez L, Gorlier A, Breard E, Sailleau C, et al. Circulation of bluetongue virus 8 in French cattle, before and after the re-emergence in 2015. Transbound Emerg Dis. 2018;65(1):281–4.

33. Breard E, Sailleau C, Quenault H, Lucas P, Viarouge C, Touzain F, et al. Complete Genome Sequence of Bluetongue Virus Serotype 8, Which Reemerged in France in August 2015. Genome announcements. 2016;4(2).

34. Rossi S, Balenghien T, Viarouge C, Faure E, Zanella G, Sailleau C, et al. Red deer (Cervus elaphus) did not play the role of maintenance host for bluetongue virus in France: The burden of proof by long-term wildlife monitoring and culicoides snapshots. Viruses. 2019;11:E903.

35. Grego E, Sossella M, Bisanzio D, Stella MC, Giordana G, Pignata L, et al. Wild ungulates as sentinel of BTV-8 infection in piedmont areas. Veterinary Microbiology. 2014;174:93–33.

36. Courtejoie N, Bournez L, Zanella G, Durand B. Quantifying bluetongue vertical transmission in French cattle from surveillance data. Veterinary Research. 2019;50:34.

37. Bournez L, Cavalerie L, Sailleau C, Bréard E, Zanella G, Servan de Almeida R, et al. Estimation of French cattle herd immunity against bluetongue serotype 8 at the time of its re-emergence in 2015. BMC Veterinary Research. 2018;14:65.

38. Nomikou K, Hughes J, Wash R, Kellam P, Breard E, Zientara S, et al. Widespread Reassortment Shapes the Evolution and Epidemiology of Bluetongue Virus following European Invasion. PLoS Pathog. 2015;11(8):e1005056.

39. Carpi G, Holmes EC, Kitchen A. The evolutionary dynamics of bluetongue virus. Journal of Molecular Evolution. 2010;70:583–92.

40. Nowak MA, Bonhoeffer S, Hill AM, Boehme R, Thomas HC, McDade H. Viral dynamics in hepatitis B virus infection. Proc Natl Acad Sci U S A. 1996;93(9):4398–402.

41. Koyuncu OO, MacGibeny MA, Enquist LW. Latent versus productive infection: the alpha herpesvirus switch. Future Virol. 2018;13(6):431–43.

42. Cohrs RJ, Gilden DH. Human herpesvirus latency. Brain Pathol. 2001;11(4):465–74.

43. Bangham CRM. Human T Cell Leukemia Virus Type 1: Persistence and Pathogenesis. Annu Rev Immunol. 2018;36:43–71.

44. Kulkarni A, Bangham CRM. HTLV-1: Regulating the Balance Between Proviral Latency and Reactivation. Front Microbiol. 2018;9:449.

45. Coffin J, Swanstrom R. HIV pathogenesis: dynamics and genetics of viral populations and infected cells. Cold Spring Harbor perspectives in medicine. 2013;3(1):a012526.

46. Gautret P, Carrara P, Parola P. Long incubation in imported human rabies. Annals of Neurology. 2014;75:324–5.

47. Cortey M, Ferretti L, Pérez-Martín E, Zhang F, de Klerk-Lorist L-M, Scott K, et al. Persistent Infection of African Buffalo (Syncerus caffer) with Foot-and-Mouth Disease Virus: Limited Viral Evolution and No Evidence of Antibody Neutralization Escape. Journal of Virology. 2019;93:e00563–19.

48. Arzt J, Fish I, Pauszek SJ, Johnson SL, Chain PS, Rai DK, et al. The evolution of a super-swarm of foot-and-mouth disease virus in cattle. PLOS ONE. 2019;14:e0210847.

49. Gray RR, Parker J, Lemey P, Salemi M, Katzourakis A, Pybus OG. The mode and tempo of hepatitis C virus evolution within and among hosts. BMC Evol Biol. 2011;11:131.

50. Osborne CJ, Mayo CE, Mullens BA, McDermott EG, Gerry AC, Reisen WK, et al. Lack of Evidence for Laboratory and Natural Vertical Transmission of Bluetongue Virus in Culicoides sonorensis (Diptera: Ceratopogonidae). J Med Entomol. 2015;52(2):274–7.

51. Nunamaker RA, Sieburth PJ, Dean VC, Wigington JG, Nunamaker CE, Mecham JO. Absence of transovarial transmission of bluetongue virus in Culicoides variipennis: immunogold labelling of bluetongue virus antigen in developing oocytes from Culicoides variipennis (Coquillett). Comp Biochem Physiol A Comp Physiol. 1990;96(1):19–31.

52. White DM, Wilson WC, Blair CD, Beaty BJ. Studies on overwintering of bluetongue viruses in insects. J Gen Virol. 2005;86(Pt 2):453–62.

53. Wilson A, Darpel K, Mellor PS. Where does bluetongue virus sleep in the winter? PLoS Biol. 2008;6(8):e210.

54. Rozo M, Gronvall GK. The reemergent 1977 H1N1 strain and the gain-of-function debate. mBio. 2015;6:e01013–15.

55. Brault AC, Powers AM, Medina G, Wang E, Kang W, Salas RA, et al. Potential Sources of the 1995 Venezuelan Equine Encephalitis Subtype IC Epidemic. Journal of Virology. 2001;75:5823–32.

56. Anderson I. Foot and mouth disease 2007: a review and lessons learned. The Stationery Office; 2008.

57. Moore SG, Hasler JF. A 100-Year Review: Reproductive technologies in dairy science. J Dairy Sci. 2017;100(12):10314–31.

58. Stevenson JS, Britt JH. A 100-Year Review: Practical female reproductive management. J Dairy Sci. 2017;100(12):10292–313.

59. Thomas FC, Singh EL, Hare WC. Control of bluetongue virus spread by embryo transfer. Progress in clinical and biological research. 1985;178:653–4.

60. Backx A, Heutink CG, van Rooij EM, van Rijn PA. Clinical signs of bluetongue virus serotype 8 infection in sheep and goats. Vet Rec. 2007;161(17):591–2.

61. Haegeman A, Vandaele L, De Leeuw I, Oliveira AP, Nauwynck H, Van Soom A, et al. Failure to remove bluetongue serotype 8 virus (BTV-8) from in vitro produced and in vivo derived bovine embryos and subsequent transmission of BTV-8 to recipient cows after embryo transfer. Frontiers in Veterinary Science. 2019;6(432).

62. Backx A, Heutink R, van Rooij E, van Rijn P. Transplacental and oral transmission of wild-type bluetongue virus serotype 8 in cattle after experimental infection. Vet Microbiol. 2009;138(3-4):235–43.

63. OIE. Terrestrial Animal Health Code. Paris: OIE; 2019.

64. Ministére de l’agriculture. Fixant les conditions sanitaires exigées pour les agréments visés à l’article L. 222-1 du code rural dans le cadre de la monte publique artificielle des animaux de l’espèce bovine. In: Journal Officiel de la Republique Francaise 2008. p. 1143.

65. Li H, Handsaker B, Wysoker A, Fennell T, Ruan J, Homer N, et al. The Sequence Alignment/Map format and SAMtools. Bioinformatics. 2009;25:2078–9.

66. Wickham H. ggplot2: Elegant Graphics for Data Analysis. 2016. Available from: https://cran.r-project.org/web/packages/ggplot2/index.html.

67. Charif D, Lobry JR. Seqin{R} 1.0-2: a contributed package to the {R} project for statistical computing devoted to biological sequences retrieval and analysis. Structural approaches to sequence evolution: Molecules, networks, populations. New York: Springer Verlag; 2007. p. 207–32.

68. Wickham H. stringr: Simple, Consistent Wrappers for Common String Operations 2019. Available from: https://cran.r-project.org/web/packages/stringr/index.html.

69. Knaus BJ, Grünwald NJ. vcfr: a package to manipulate and visualize variant call format data in R. Molecular Ecology Resources. 2017;17:44–53.

70. Magoč T, Salzberg SL. FLASH: Fast length adjustment of short reads to improve genome assemblies. Bioinformatics. 2011;27:2957–63.

71. Langmead B, Salzberg SL. Fast gapped-read alignment with Bowtie 2. Nature Methods. 2012;9:357–9.

72. Roberts A, Pachter L. Streaming fragment assignment for real-time analysis of sequencing experiments. Nature Methods. 2013;10:71–3.

73. Wilm A, Aw PPK, Bertrand D, Yeo GHT, Ong SH, Wong CH, et al. LoFreq: A sequence-quality aware, ultra-sensitive variant caller for uncovering cell-population heterogeneity from high-throughput sequencing datasets. Nucleic Acids Research. 2012;40:11189–201.

74. Nagarajan N, Kingsford C. GiRaF: Robust, computational identification of influenza reassortments via graph mining. Nucleic Acids Research. 2011;39:e34.

75. Huelsenbeck JP, Ronquist F. MRBAYES: Bayesian inference of phylogenetic trees. Bioinformatics. 2001;17(8):754–5.

76. Guindon S, Dufayard JF, Lefort V, Anisimova M, Hordijk W, Gascuel O. New algorithms and methods to estimate maximum-likelihood phylogenies: Assessing the performance of PhyML 3.0. Systematic Biology. 2010;59:307–21.

77. Suchard MA, Lemey P, Baele G, Ayres DL, Drummond AJ, Rambaut A. Bayesian phylogenetic and phylodynamic data integration using BEAST 1.10. Virus Evolution. 2018;4:vey016.

78. Guindon S, Delsuc F, Dufayard JF, Gascuel O. Estimating maximum likelihood phylogenies with PhyML. Methods Mol Biol. 2009;537:113–37.

79. Rambaut A, Lam TT, Max Carvalho L, Pybus OG. Exploring the temporal structure of heterochronous sequences using TempEst (formerly Path-O-Gen). Virus Evolution. 2016;2:vew007.

80. Drummond AJ, Ho SYW, Phillips MJ, Rambaut A. Relaxed phylogenetics and dating with confidence. PLoS Biology. 2006;4:699–710.

81. Posada D. jModelTest: phylogenetic model averaging. Mol Biol Evol. 2008;25(7):1253–6.

82. Minin VN, Bloomquist EW, Suchard MA. Smooth skyride through a rough skyline: Bayesian coalescent-based inference of population dynamics. Molecular Biology and Evolution. 2008;25:1459–71.

83. Bürkner PC. brms: An R package for Bayesian multilevel models using Stan. Journal of Statistical Software. 2017;80:1–28.

84. Yu G, Smith DK, Zhu H, Guan Y, Lam TTY. ggtree: an r package for visualization and annotation of phylogenetic trees with their covariates and other associated data. Methods in Ecology and Evolution. 2017;8:28–36.

85. Arnold JB. ggthemes: Extra Themes, Scales and Geoms for ‘ggplot2’ 2019. Available from: https://cran.r-project.org/web/packages/ggthemes/index.html.

86. Wilke CO. cowplot: Streamlined Plot Theme and Plot Annotations for ‘ggplot2’ 2019. Available from: https://cran.r-project.org/web/packages/cowplot/index.html.

87. Kahle D, Wickham H. ggmap: Spatial visualization with ggplot2. The R Journal. 2013;5:144–61.

88. Garnier S. viridis: Default Color Maps from ‘matplotlib’ 2018. Available from: https://cran.r-project.org/web/packages/viridis/index.html.

89. Kay M. tidybayes: Tidy Data and Geoms for Bayesian Models 2019. Available from: https://cran.r-project.org/web/packages/tidybayes/index.html.

90. Grolemund G, Wickham H. Dates and times made easy with lubridate. Journal of Statistical Software. 2011;40.

91. Pebesma EJ, Bivand RS. Classes and methods for spatial data in {R}. R News. 2005;5.

92. Hijmans RJ. raster: Geographic Data Analysis and Modeling 2019. Available from: https://cran.r-project.org/web/packages/raster/index.html.

93. Bivand RS, Lewin-Koh N. maptools: Tools for Handling Spatial Objects 2019. Available from: https://cran.r-project.org/web/packages/maptools/index.html.

94. Bivand RS, Rundel C. rgeos: Interface to Geometry Engine - Open Source (‘GEOS’) 2019. Available from: https://cran.r-project.org/web/packages/rgeos/index.html.

95. Bivand RS, Keitt T, Rowlingson B. rgdal: Bindings for the ‘Geospatial’ Data Abstraction Library 2019. Available from: https://cran.r-project.org/web/packages/rgdal/index.html.

96. Pebesma EJ. Simple features for R: Standardized support for spatial vector data. The R Journal. 2018;10:439–46.

97. Schnute JT, Boers N, Haigh R. PBSmapping: Mapping Fisheries Data and Spatial Analysis Tools 2019. Available from: https://cran.r-project.org/web/packages/PBSmapping/index.html.

